# Stelar starch management tailors diurnal and rehydration-related water flows in *Pinus pinea* needles

**DOI:** 10.64898/2026.03.06.710090

**Authors:** Peter Bork, Chen Gao, Emilie Thor Herfelt, Margaux Schmeltz, Tomas Bohr, Alexander Schulz

**Author notes:** Contributed equally to this work (Peter Bork, Chen Gao). E-mail addresses.

## Abstract

Pine needles contain two vascular cell types unique to gymnosperms: Transfusion parenchyma (tp) and tracheids (tt). Since they form the only connections between vascular bundles and bundle sheath, we hypothesised that they are involved in regulating the needle’s water import and photoassimilate export. Synchrotron-based tomography enabled us to quantify volume changes of tp and tt cells in *Pinus pinea* needles systematically along the needle and throughout a diurnal day cycle, as well as under rehydration. As a physiological indicator of tp’s carbohydrate status served their starch content. Segmentation of the comprehensive data uncovered dramatic volume changes during dehydration and showed a diurnal course of starch formation and degradation. These changes suggest a yet unknown osmotic water flux between tp and tt, balanced by the former’s carbohydrate status. Confirming our hypothesis, excess of photoassimilates in tp cells went into starch synthesis during the day. Starch mobilisation during the night increased the osmotic potential in tp and led to water intake. According to the decreasing starch fraction from base to needle tip, this mechanism is predominant in the upper needle segments, particularly after rehydration of dehydrated needles. Mechanistically, osmolytes in tp cells maintain tension in tt for the needle’s water import.

**Highlight:** Synchrotron tomographic microscopy uncovers diurnal starch fluctuations and osmotic water pumping in inner tissues of pine needles that are utilised at night and when recovering from dehydration

## Introduction

In the needle-shaped leaves of conifers, a highly specialised tissue mediates post-xylem nutrient and pre-phloem assimilate transport between the axial vascular tissues and the radially organised mesophyll. This transfusion tissue consists of living transfusion parenchyma cells (tp) and a system of dead transfusion tracheids (tt) that extends the xylary apoplast (Hacke *et al*., 2015; Hu and Yao, 2011; Huber, 1947). The transfusion tissue is bordered by the endodermis-type bundle sheath (hereafter referred to as ‘endodermis’ for simplicity). The endodermis forms a barrier between the vascular and the cortical apoplast of the mesophyll. Accordingly, the outward flow of water and nutrients meets the inward symplasmic flow of photoassimilates in the endodermis (en) cells (Mai *et al*., 2024). The bulging shape of tp cells indicates a positive turgor pressure, osmotically generated by the solute potential of the tp cells’ cytoplasm, directly bordering the water-storing tt system.

Earlier work (Gao et al. 2023) showed an approximately 2:1 volume proportion of living tp to dead tt across different Pinaceae. We wondered, whether this proportion is maintained constant over the day, and hypothesised that tp cells control the cytoplasmic concentration of photoassimilates by a homeostatic mechanism. This mechanism could for instance involve the mutual exchange of osmotically effective sugars with osmotically ineffective starch. Sequestering soluble carbohydrates in starch is a means to control the cytoplasmic solute potential in tree stems (Furze *et al*., 2018).

Such a regulatory mechanism might be key for cavitation repair or specific environmental conditions such as freeze-thaw cycles where the entire needle is dehydrated (Bouche *et al*., 2016; Mayr *et al*., 2014). Both situations were shown to lead to deformation of transfusion tissue and, in the case of desiccation-tolerant ferns, to a rehydration of living vascular cells before xylem element refilling (Brodribb and Holbrook, 2005; Hacke *et al*., 2015; Holmlund *et al*., 2019).

We chose to study diurnal changes in the transfusion tissue and the response to dehydration and rehydration by live imaging of pine needles at the Tomcat beamline of the Poul Scherrer Institute (PSL) in Villigen, Switzerland. Here, we could apply tomographic microscopy on pine needles with a sufficient high temporal and spatial resolution. Using propagation-based phase contrast, the different cell types of the vascular tissue could be identified, and even subcellular details were resolved, such as starch grains (Mai *et al*., 2024). In contrast to µXCT from a lab X-ray source, identification of tissues did not depend on contrast agents as in (Gao *et al*., 2023).

In angiosperm leaves, mesophyll chloroplasts contain transitory starch that follows diurnal variations. The size of transitory starch grains depends on the balance between photosynthetic sugar formation and sugar export trough the phloem. Typically, the highest starch amount is found in the afternoon which is totally mobilised through the night and allows the leaf to maintain phloem export and respond to abiotic and biotic stress (Gersony *et al*., 2020; Ribeiro *et al*., 2022; Tixier *et al*., 2018). When phloem export was inhibited by antisense repression of phloem loading, large starch grains were formed in all mesophyll cells (Schulz *et al*., 1998). Conifer needles use a symplasmic phloem loading strategy as suggested by a continuous symplasmic pre-phloem pathway and confirmed by phloem tracer experiments (Gao *et al*., 2023; Han *et al*., 2022; Liesche *et al*., 2011; Liesche and Schulz, 2012; Liesche and Schulz, 2018). The pre-phloem pathway starts in the endodermis layer and continues via a few tp cells to the phloem flank, where Strasburger cells contact the axial phloem sieve elements. As angiosperm leaves, they also seem to have diurnal changes in carbohydrate content (Liesche *et al*., 2021; Webb and Kilpatrick, 1993). Parameters which might determine the diurnal assimilate stream in the pre-phloem pathway are photosynthesis or mobilisation of transitory starch in the mesophyll, the assimilate gradient from endodermis to phloem, mobilisation of en and tp starch and the phloem export rate, respectively the assimilate concentration in the axial phloem. These parameters have influence on the cell turgor in the assimilate transporting cells (Ali *et al*., 2023). However, it is not known whether the starch biochemically detected at different timepoints and positions in the needle originated from mesophyll chloroplasts and/or from the transfusion tissue. Transitional starch in tp and en cells was recently discussed to play an important role in maintaining a steady carbohydrate supply for phloem loading, especially during periods of hight transpiration (Vollenweider *et al*., 2026). As a drought-tolerant, Mediterranean pine species we chose *P. pinea* and scanned the needle segments at three positions: the tip, the mid and the base to take into account possible physiological differences between these positions, such as those related to starch metabolism (Liesche *et al*., 2021).

## Material and Methods

### Plant material

For electron microscopy, *Pinus pinea* L. branches were collected from a 4-yr-old tree in the glasshouse of the University of Copenhagen, Denmark, and, for tomographic microscopy, from a tree in the old Botanical Garden of the University of Zürich. Up to scanning, these branches were kept hydrated in a cabinet at room temperature with a 12/12 h day/night photoperiod, starting illumination with a 60-Watt glow lamp at 7 a.m. Needles taken for our tomographic scans in March 2023 were last-year (2022) needles. For assessing changes during a diurnal day cycle, we scanned the needles at five timepoints, three during the day and two during the night. For the dehydration and rehydration experiment, needles were detached from the branch and left to accommodate to dim room light on a table for 24 hours. In order to compare needles of a similar diurnal stage, control and dehydrated needles were scanned between 6 p.m. and midnight. Rehydration was done by immersing the needle base 10 mm deep in Eppendorf tubes filled with water and kept there for 6 hours.

### Electron microscopy

After gently removing the epidermis with sandpaper, 10-mm needle segments were immediately immersed into Karnovsky’s fixative (4% (w/v) paraformaldehyde and 5% (w/v) glutaraldehyde in 0.1M sodium cacodylate buffer, pH 7.4), and fixed according to (Hunziker and Schulz, 2019). To avoid preparation artefacts, 3-mm ends of each segment were discarded. After polymerization, ultrathin sections 70 nm thickness were cut with a diamond knife and ultramicrotome (EM UC7; Leica Microsystems, Wetzlar, Germany). The sections were transferred to film-coated single-slot grids (FCF2010-CU; Electron Microscopy Sciences, Hatfield, PA, USA) and post-contrasted with uranyl-less solution for 3 min and lead citrate solution for 3 min with thorough washing after each step. For transmission electron microscopy (TEM), a ThermoFisher Talos L120C G2 was used at 120 kV acceleration voltage and stitched using MAP 3 software where necessary.

### Tomographic microscopy at the TOMCAT beamline, Paul Scherrer Institute, Switzerland

Needles were detached from the branch under water and immediately transferred to 2 ml Eppendorf tubes filled with tap water. After detachment each needle base was placed in water-filled Eppendorf tubes, and each needle was scanned at three positions, 10 mm from the tip, in the mid (80 mm from the tip) and at the base (10 mm from the detachment site), starting with the mid segment; for the tip, the needle was cut and placed in water at the mid scanning site, and, finally, the removed basal needle part was inverted, placed in water and subsequently scanned. Scanning of each 0.7 mm segments took some 5 min, and the scanning of one needle at the three positions in average 20 minutes, preparation time included. We used for all measurements three biological replicates and, as technical replicates, three segmentations of each location at the cross-sectional frame numbers 500, 1000 and 1500 (of 2160 frames per 0.7 mm segment). The needles were held upright during rotation in a home-made chamber while immersed in the Eppendorf tube (Gao et al., 2023).

For imaging, the middle, tip, and base segment were targeted, with 1000 projections acquired at 21 keV over 180° rotation around the needle centre using a × 20 objective, resulting in an effective pixel size of 0.325 μm. Reconstruction of the scans followed the gridrec and paganing algorithms established at the TOMCAT beamline, providing z-stacks of 2160 needle cross sections. To avoid optical artefacts at the ends of the scanned area, 2D-segmentations were taken at cross section 500, 1000 and 1500. The dimensions of the scanned image cube were c. 0.7 × 0.7 × 0.7 mm. To visualize the 3D structure of the interior of the needle, we imported an 350 µm high z-stack (1000 cross sections) into the open-source software FLUORENDER (https://www.sci.utah.edu/software/fluorender.html), see Supplementary video.

### Segmentation

Avizo software were used to segment from each needle the tip, mid and base segment and from each of them position 500, 1000, and 1500 of the respective image stacks. Needles have two vascular bundles inside endodermis-bordered stele. With the chosen magnification, only half of the stele was covered within the field of view. This half was selected, segmenting the endodermis (en), transfusion parenchyma (tp) and the transfusion tracheid space (tt). Recognition of the tissue types was based on the difference in X-ray contrast, which was assigned to five increasing X-ray densities: air, water-filled axial xylem and tt, cytoplasm, cell walls and lignified cell walls of the xylem or starch grains, visualised here as five grey values from dark to bright. Starting with marking the space between endodermis and vascular bundle with the brush tool, all tp cells, the bordering en cells and all undefined areas were subtracted from the initial space and defined as tt space. Starch grains in tp and en cells were identified, in doubt by checking the neighbouring cross-sectional slices, and marked in the selected slice position. After using Avizo’s statistics function, the segmentations were exported as csv. These files contained specimen numbers, the cross-sectional areas of the selected cell types, scanning time point, numbers and area of starch grains per cell and cellular starch fraction.

### Data analysis

Microsoft Excel software was used for filtering the respective cell types, their areas and the starch area, respectively the cellular starch fraction, and transferred into the Prism software, where the statistic tests were run and the graphs created. Significance of differences between data points was tested with Anova, is shown with letters above the violin plots and defined in the legends. The solid line in violin plots defines the median; the stippled lines the 25 % percentiles.

## Results

Reconstruction of transverse needle sections from phase-contrast tomography series enabled us to discriminate the axial vascular tissues, the transfusion tissue with transfusion parenchyma and transfusion tracheids and the endodermis surrounded by the mesophyll (Fig. 1A). A curving longitudinal section just inside the endodermis confirmed that the only two cell types bridging the distance between endodermis and the axial vascular tissues are tp and tt (Fig. 1B). All tt-cells of a cross section were segmented as one system as they physiologically behave as a continuous extension of the axial xylem; see (Gao *et al*., 2023). Contrast differences were utilised for segmentation of the en-bordered transfusion tissue, considering and quantifying only half of the stele in this paper (compare Fig. 1C, D: tp and en cells are shown in different colours, the tt system in magenta). Starch grains were detected in tp and en cells; their expectable presence in the mesophyll was masked by the phase-contrast induced artefactual halos around living cells bordering the intercellular airspace. TEM images through the stelar tissue confirmed that the bright spots detected in synchrotron images indeed were starch grains and identified tp cells by prominent tannin vacuoles and en cells by small spherical vacuolar inclusions (Fig. 1E, F). By size and shape the starch grains in tp and en cells are distinct from the flat and discoidal transitory starch grains in chloroplasts of herbaceous plants depicted in textbooks (cf. (Smith and Zeeman, 2020).

**Figure 1:**
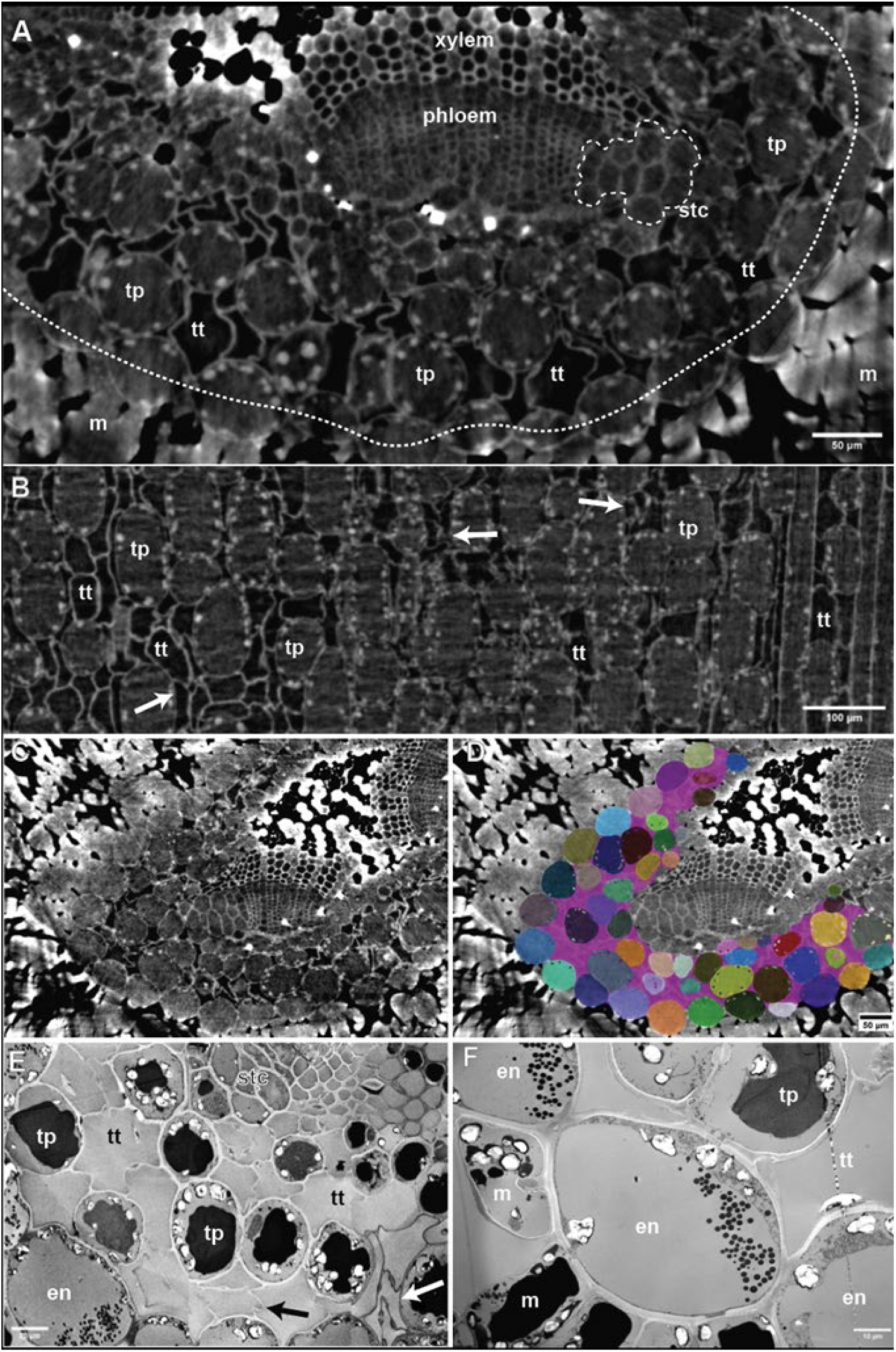
X-ray and electron micrographs of vascular tissues in *Pinus pinea* needles. detached immediately before imaging (A-E). **A:** Representative cross section through a needle at the base, reconstructed from a tomographic synchrotron series. The transfusion tissue is placed between endodermis (en) and the axial xylem (ax) and phloem (ap). Large starch grains (white arrows) occurred in the periphery of tp and en cells. **B:** Curving longitudinal section inside the endodermis, comprising half the stele. Only en and tt cells are found. Arrows show folded tt cell walls **C and D:** Segmentation enabled quantification of the number and cross-sectional area of cell types and of their starch grains, respectively. Marked in magenta is the entire continuous system of tt cells. **E:** Electron micrograph from the flank of one of the two vascular bundles, containing Strasburger cells (stc), tp cells with peripheral starch grains and tannic acid-filled central vacuoles. Both, tp and tt cells are in contact with en cells that are recognized by small circular tannic-acid vesicles and contain starch grains at their periphery (arrows). In **(F)**, also some mesophyll cells (ms) are depicted with chloroplasts, containing transitory starch, and tannic-acid-filled central vacuoles.

### Structural differences along the needle axis

Confirming the structural data from (Gao *et al*., 2023), the diameter of the stele increases from tip to base as does the number of axial phloem and xylem elements (for representative examples see Supplementary Figure S1). The resolution power of phase-contrast tomographic microscopy enabled us to visualise starch grains in endodermis and stele of intact needles. Biochemically, these starches are difficult to discriminate from mesophyll starch (but see upcoming methods by (Vollenweider *et al*., 2026). First, we asked whether the distribution of tp and en starch is even along the needle axis, or whether there is a gradient in the starch content. As a physiologically relevant measure of starch amounts in cells, we chose the starch fraction (starch area per cell area). Figure 2A combines all recordings taken during 24 hours of this synchrotron experiment. In both tissues, tp and en, the starch fraction was smallest in the tip and about double that in the base. In the mid segment, the starch fraction in tp cells was in between tip and base, while that in en cells equalled the tip. Tp cells contained a roughly more than 50 % larger starch fractions than endodermis cells in all positions (Figure 2A). This difference seems partially to be due to the general difference in cell size, since the cross-sectional area of en cells is between 22 and 49 % larger than that of tp cells. Overall, the starch content increased from tip to base and was larger in tp than en cells whether quantified as fractional area, grain size, or grain number (Table 1).

**Figure 2:**
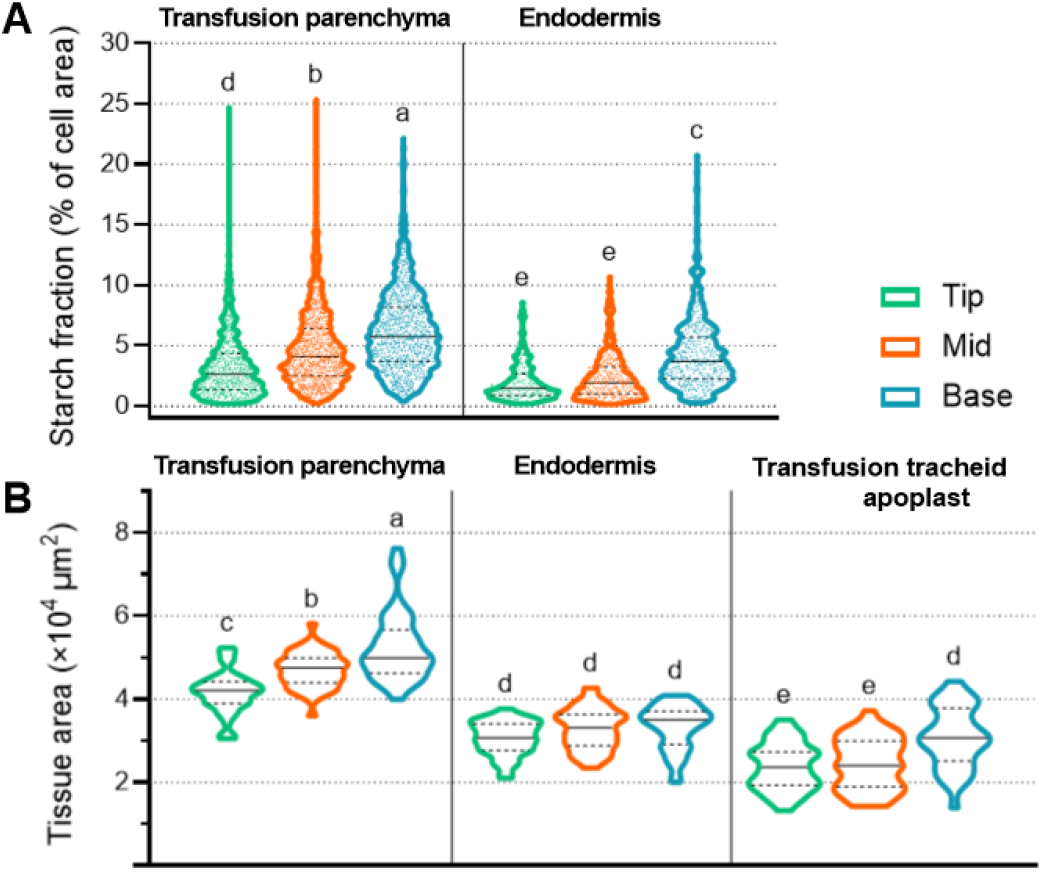
Starch fraction and tissue area differences along the needle axis of *P. pinea* as revealed in tomographic microscopy series taken throughout the day **A:** Axial starch gradient in needles and quantified as fraction of the cross-sectional area of transfusion parenchyma and endodermis cells. Both tissues accumulate more than two times more starch in the base than in the tip segment (n>837 and >261 per axial location for tp and en, respectively; different letters mark significant differences (P<0.0001) according to one-way ANOVA followed by Tukey’s post-hoc test. **B:** Cross-sectional tissue areas. The tp tissue area increases from tip to base. There is no significant difference in the area of the endodermis along the axis. The apoplast space formed by the tt system is significantly larger in the base segments than in mid and tip (n>39). Different letters mark significant differences (*P*<0004) according to one-way ANOVA followed by Tukey’s post-hoc test.

**Table 1.**
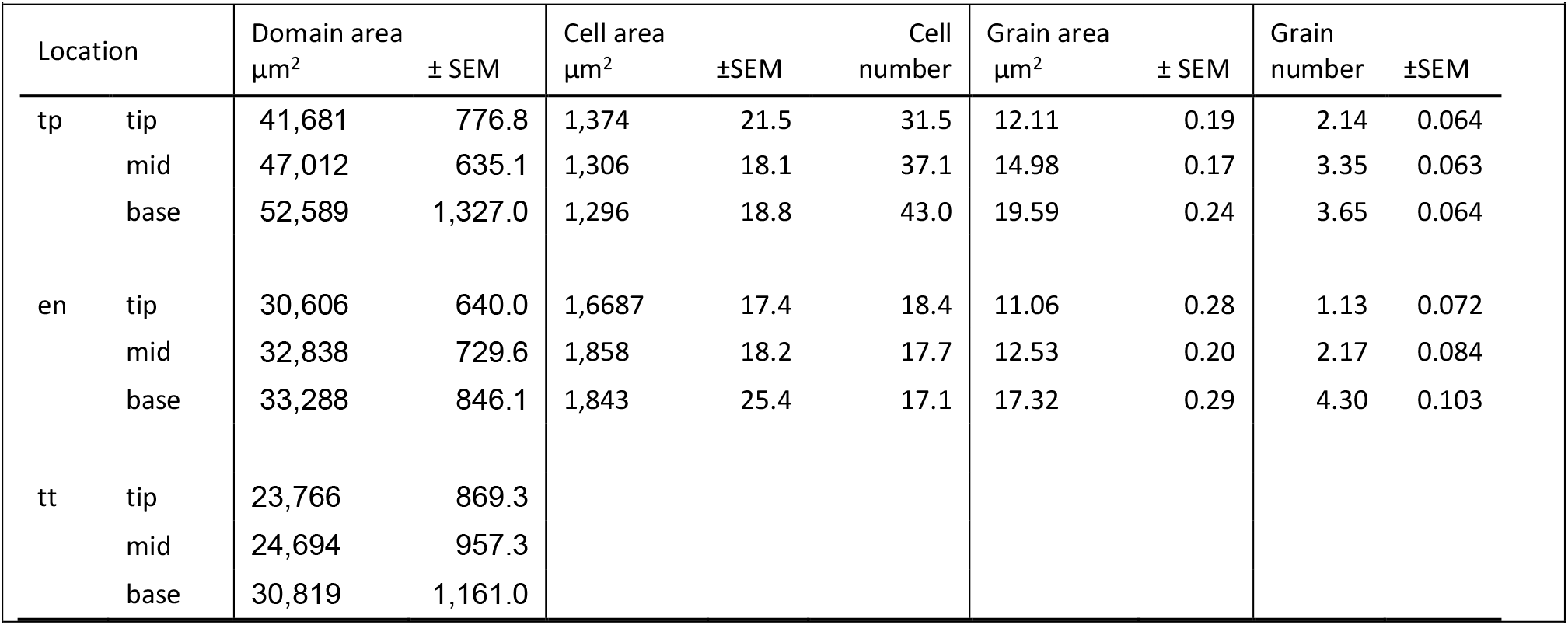
Mean domain area, cell area and number, and mean starch area and number.

|  |  |  |  |  |  |  |  |  |  |  |
| --- | --- | --- | --- | --- | --- | --- | --- | --- | --- | --- |
| Location | | Domain area<br>$\mu\text{m}^2$ ± SEM | | Cell area<br>$\mu\text{m}^2$ ±SEM | | Cell<br>number | Grain area<br>$\mu\text{m}^2$ ± SEM | | Grain<br>number | ±SEM |
| tp | tip | 41,681 | 776.8 | 1,374 | 21.5 | 31.5 | 12.11 | 0.19 | 2.14 | 0.064 |
|  | mid | 47,012 | 635.1 | 1,306 | 18.1 | 37.1 | 14.98 | 0.17 | 3.35 | 0.063 |
|  | base | 52,589 | 1,327.0 | 1,296 | 18.8 | 43.0 | 19.59 | 0.24 | 3.65 | 0.064 |
| en | tip | 30,606 | 640.0 | 1,6687 | 17.4 | 18.4 | 11.06 | 0.28 | 1.13 | 0.072 |
|  | mid | 32,838 | 729.6 | 1,858 | 18.2 | 17.7 | 12.53 | 0.20 | 2.17 | 0.084 |
|  | base | 33,288 | 846.1 | 1,843 | 25.4 | 17.1 | 17.32 | 0.29 | 4.30 | 0.103 |
| tt | tip | 23,766 | 869.3 |  |  |  |  |  |  |  |
|  | mid | 24,694 | 957.3 |  |  |  |  |  |  |  |
|  | base | 30,819 | 1,161.0 |  |  |  |  |  |  |  |

Figure 2B shows that the stelar area taken by tp cells increases from tip to base significantly with 26 % (mean from 41,700 to 52,600µm^2^ for half the stele, see Table 1). This reflects an increase of the number of tp cells; their cross-sectional area was constant at around 1300 µm^2^. By contrast, the endodermis area stayed at 31-33,000 µm^2^ with around 18 cells per half stele. The tt system did not change much between tip and mid position but expanded with 30 % in the base (Fig. 2B; Table 1). Considering the increase in both, cell number and starch fraction, the tp tissue accumulated proportionally more starch in the mid and base position than the en layer.

### Diurnal changes of transfusion tissue and endodermis

To assess diurnal changes, branches with last-year needles were kept at a 12/12 h day/night photoperiod and detached from the branch immediately before scanning. In tp cells, the starch fraction increased continuously over the day up to the afternoon but dropped significantly to their smallest values at night, i.e. at the midnight and early morning values (Fig. 3A). In en cells, the starch fraction showed a similar diurnal course with the smallest values in the early morning (Fig. 3B). To assess the dynamic range of starch grain synthesis and degradation, we analysed the frequency distributions of cross-sectional grain areas. These showed that the grains were smallest at night, and significantly larger at daytimes, amounting to an increase of some 20% for tp starch at each of the three needle positions and a similar difference for en starch except in the tip (Figure 3C,D). The size range of grains spanned 2.8 µm^2^ (detection limit) to 60 µm^2^; Gaussian fit centres showed that the largest differences in the tip segment equals an increase in the starch grain area of 53 % (8.81 and 13.51 µm^2^ for the last night and last day value, respectively; for details of the diurnal changes of starch fraction and grain size, see Supplementary Figure S2). Assuming spherical grain shapes, 53 % increase in area translates to 89 % increase in starch volume.

**Figure 3:**
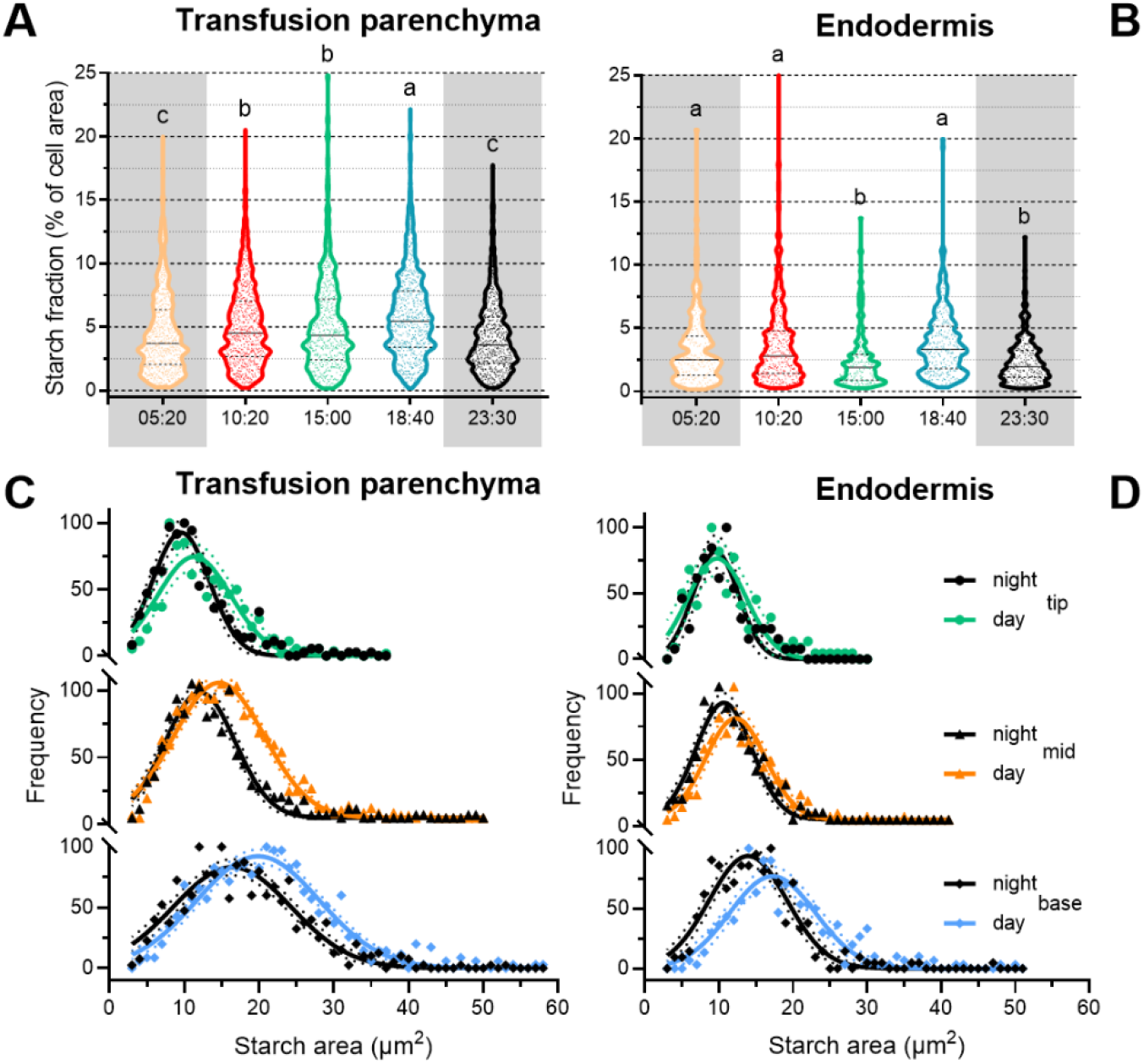
Diurnal starch changes in transfusion parenchyma and endodermis. **A:** Starch fraction of tp cross-sectional cell areas integrated over tip, mid and base segments of tomographic microscopy series, each segmented at three positions (n>550 per time point). The starch fraction of tp cells increases up to the evening and is smallest at the midnight value. Different letters mark significant differences (*P*<0.005 according to one-way ANOVA followed by Tukey’s post-hoc test). **B:** Starch fraction of en cross-sectional cell areas integrated over tip, mid and base segments of µX-ray CT series, each segmented at three positions (n>189 per time point). The only significant difference lies between a high evening value and early morning (*P*<0.0001 according to one-way ANOVA followed by Tukey’s post-hoc test). **C and D:** Normalized frequency distributions of starch grain sizes in tip, mid and base needle segments for tp and en, respectively. In all segments, the grains were smaller a night than at daytimes. Curve fittings (lines) assume a Gauss distribution of grain sizes and show significant differences between night and day curves in each needle segment for tp starch according to 95 % confidence intervals. For en starch, the curves in the tip and mid segment were overlapping. Confidence intervals at the 95% level are indicated.

The observed diurnal starch changes indicate that starch grains increased in size during the day and were mobilised during the night like transitory starch in the mesophyll of most plants. In contrast to transitory starch, however, we did not observe a complete disappearance of tp starch at the end of night. Hypothesising that the partial mobilisation of starch would lead to formation of mono- and disaccharides, and thus to an increase in turgor and concomitant cell volume changes, we measured the cross-sectional area of the starch-containing cells (tp and en) and that of the tt apoplast domain. Indeed, the absolute cell areas of tp and en changed moderately throughout the day and were smallest at the 15-o’clock value to increase to the midnight timepoint, after having a first peak at the first day timepoint (Fig. 4A, Supplementary Fig. S3). By contrast, the area of the apoplasmic tt-system showed strong significant changes, amounting to a 50% increase from the last day to midnight value (Fig. 4A). In contrast to the cellulosic cell walls of tp and en cells, the thin lignified cell walls subdividing the tt space can be found heavily folded (Fig. 1B and E: arrows) enabling stronger volume reductions than turgid walls of living cells. To test the possibility that cell sizes are biased by size variations between individual needles, we measured the absolute area, taken together by the en layer, the tp tissue and the apoplasmic tt system. Supplementary Figure S4A shows that there was generally little diurnal variation of the summed area, allowing to reject a bias through natural variation. However, the midnight values stood out, being significantly larger than at the neighbouring timepoints. Obviously, the swelling of the entire stele was dominated by the apoplasmic tt space, while the areas of tp and en cells were constant along the needle (Table 1).

**Figure 4:**
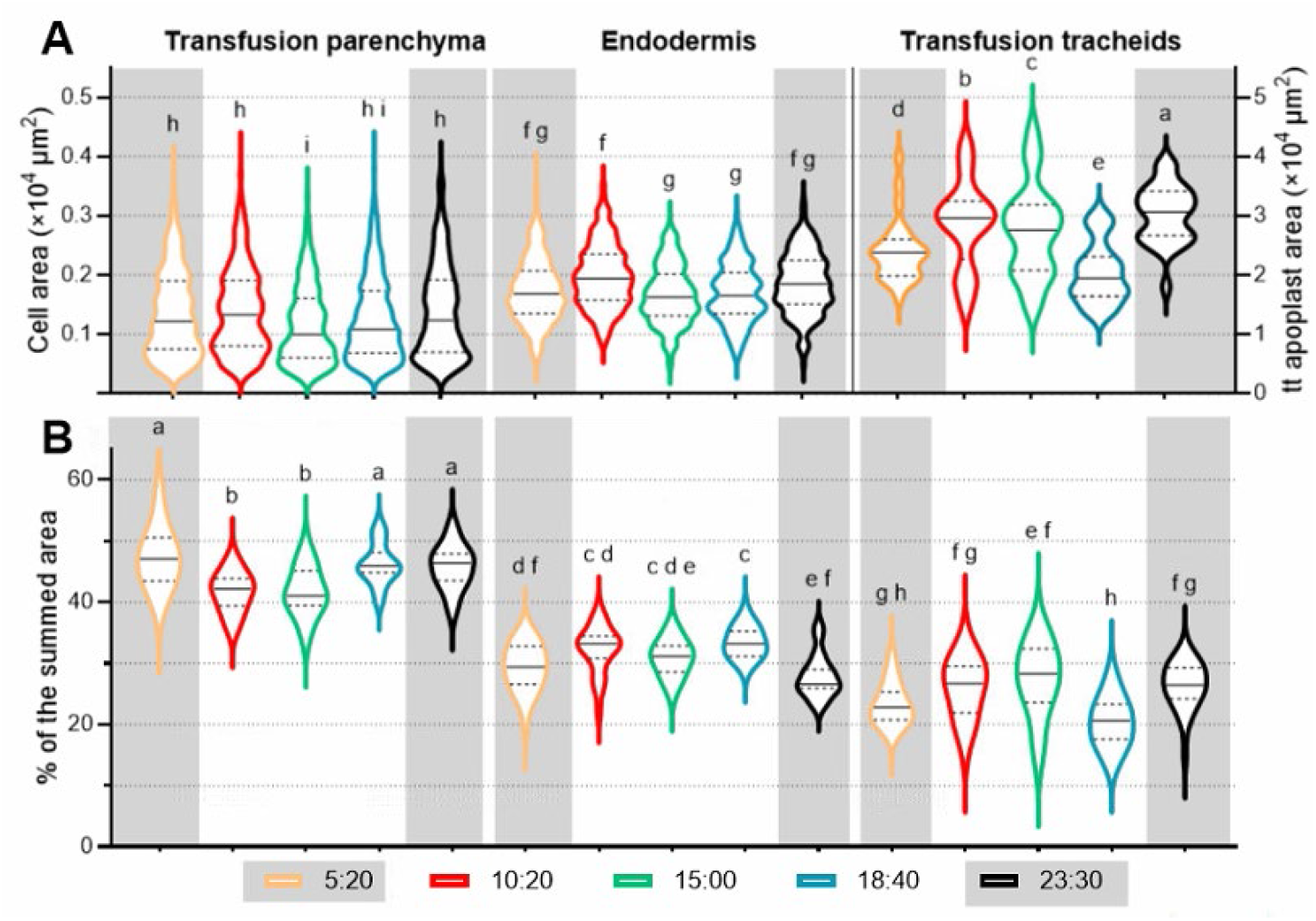
Diurnal tissue area changes in the needle. **A:** The cross-sectional cell area of tp cells ranged from 100 to 4000 µm^2^ integrated over tip, mid and base positions of tomographic microscopy series, while that of en cells from 420 to 3700 µm^2^ (n>751 for tp; >385 for en). Cell area increased in both tissue types from afternoon timepoint to midnight. En cells showed also an increase between the last night and the morning timepoint. For changes of the cell areas in each of the positions see supplementary Figure S3. Individual transfusion tracheids (tt) could not be segmented; the entire tt system ranged from 13000 to 44000 µm^2^ per cross section (n>21) The smallest and largest sizes are between last day and first nighttime value. Different letters mark significant differences between neighbouring time points (P<0.0001 according to ANOVA followed by Tukey’s post-hoc test). **B:** The relative distribution of cross-sectional areas. While the tp domain decreased up to the afternoon and increased thereafter, the apoplast of the tt system increased first to drop in the evening and increase thereafter again. The en layer did not seem to be correlated with either tp or tt [n> 21 per time point that relates the sum of tp cell areas, sum of en cell areas and the area of the tt system to the summed area (tp + en + tt); P<0.0001 according to ANOVA followed by Tukey’s post-hoc test].

To get a realistic picture of osmotic volume changes between the living cells (tp and en) and the stellar apoplast (tt system), we quantified the diurnal variations of the proportion, each of these spaces was inhabiting in the summed area (tp + en + tt). Figure 4B shows that the tp domain was smallest during the day and achieved a significantly larger part of the summed area in the evening and during the night. By contrast, the area taken by the tt apoplast was smallest in the early morning and largest at noon and midnight, mirroring tp’s volume changes except for the midnight timepoint. Here, the tt apoplast area increased in volume, apparently taking volume released from the en cell layer. The space taken by the en layer showed less fluctuations than tp or tt and did not follow either’s course (Fig. 4B). The opposite fluctuations in tt and tp volumes over an entire day (Fig. 4B) support the hypothesis that water osmotically moves in and out of tp cells depending on the water potential difference between these two tissues. Aquaporins have been shown in tp and en plasma membranes, both of which contact the tt space directly (Laur and Hacke, 2014).

The observed changes were significant, but relatively small. To understand the role of the transfusion tissue in more extreme situations, we subjected needles to dehydration aiming at diminishing the water storing capacity of the tt and, thus, changing the osmotic balance in the transfusion tissue (and possibly also in the en cell layer).

### Dehydration-related changes of transfusion tissue and endodermis

Segmentation of dehydrated and rehydrated needles showed dramatic changes in the transfusion tissue. Representative segmentations of (A) control, (B) dehydrated and (C) rehydrated needle cross sections are shown in Figure 5. Twenty-four hours after detachment of a needle, the tt system has shrivelled up but resumed fully when the needle base was immersed in water (Fig. 5A-C: see the magenta marked tt system). The dehydrated apoplast domain of the tt system was down to 39 % of the control value (Fig. 5D). Not only did the tt system respond to dehydration, but also the entire inner needle area (tt + tp + en) was significantly reduced (Supplementary Figure S4B). In fact, dehydration-related water exosmosis from tp and en cells led to significant reduction of their cell sizes (Fig. 5D).

**Figure 5:**
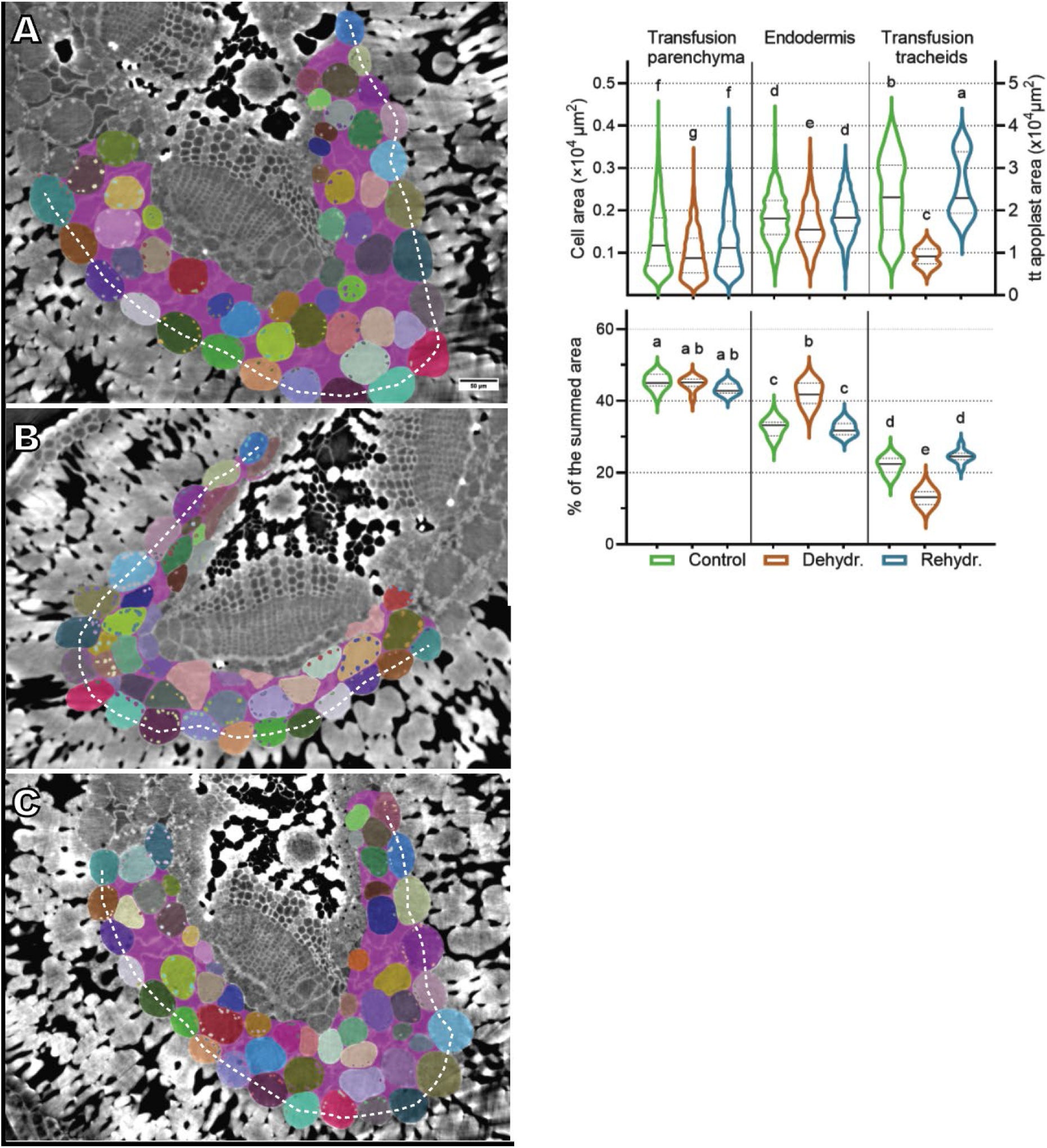
Dehydration-related tissue area changes in transfusion tissue and endodermis. **A-C**: Segmented cross-sectional examples of control (**A**), dehydration (**B)** and rehydration (**C**). Each segmented cell is coloured, the endodermis moreover labelled with a stippled line; the tt system is shown in magenta. **D, E**: Quantification of area changes in tomographic microscopy series, integrated over tip, mid and base segment. **D:** The cross-sectional areas of tp and en cells dropped under dehydration and resumed after rehydration. For differences along the needle axis see Supplementary Fig. S5. The tt space shrunk dramatically under dehydration but recovered completely after rehydration. Different letters assign significant differences (P<0.0001 according to ANOVA followed by Tukey’s post-hoc test); tp and en n>458, tt space n >27). **E:** The relative distribution of cross-sectional areas. Dehydration and rehydration did not change the proportion of the tp domain, while the tt apoplast domain strongly dropped under dehydration and resumed fully when rehydrated. The en layer behaved opposite to tt (n> 9 per treatment that relates the sum of tp cell areas, sum of en cell areas and the area of the tt system to the summed (tp+en+tt) in the respective cross section. Different letters mark significant differences; between tp and tt or en; P<0.0001 according to ANOVA followed by Tukey’s post-hoc test)).

When quantifying the proportions between tt, tp and en of the summed area, the tp domain kept its proportion (some 42 %) independent of the treatment, while the apoplast of the tt system dropped from 22 to 13 % under dehydration, and the en layer increased its proportion (Fig. 5E). Six hours of rehydration let the needles resume their tissue proportions and all three tissues returned to control values. Resumption of needle architecture and tissue proportions confirms that the studied needles, despite 24-h detachment, can cope with imminent air bubble formation of the xylem.

The observed relative and absolute changes support the idea that the tt system is a water-storing, sponge-like tissue (Gao *et al*., 2023) that can accommodate large changes in water potential by shrinking and swelling. This raised the question whether refilling of the apoplasmic tt-space resulted from the specific structural properties of tt cell walls or rather involved active responses of living needle tissue. Such responses might be postulated considering the natural adaptive range of needles that also on tree are exposed to climatic challenges like drought and freezing.

We checked the size of starch grain in tp and en cells after detachment to detect indications of enzymatic degradation activities. Figure 6A shows that dehydration followed by rehydration reduced the starch fraction in the tp domain, while that of en cells was constant. However, increase of the starch fraction in en cells under dehydration is biased by the shrinking of the cell volume (see Fig. 5). A more reliable indication of enzymatic activity, necessary if starch should be mobilised in tp and en cells, are reductions in the cross-sectional starch area during the experiment. Indeed, significantly smaller grains than in controls occurred after rehydration in tp and en cells of all needle positions according to their Gaussian fits of their frequency distribution (Fig. 6B). In the tip segment, they amounted to >40 % reduction in grain area (both tp and en) and >20 % reduction in mid and base segment for tp cells (>11 % for en cells). Assuming spherical grains in tp and en cells (Fig. 1), 42 % reduction in area translates to 56 % reduction of starch volume.

**Figure 6:**
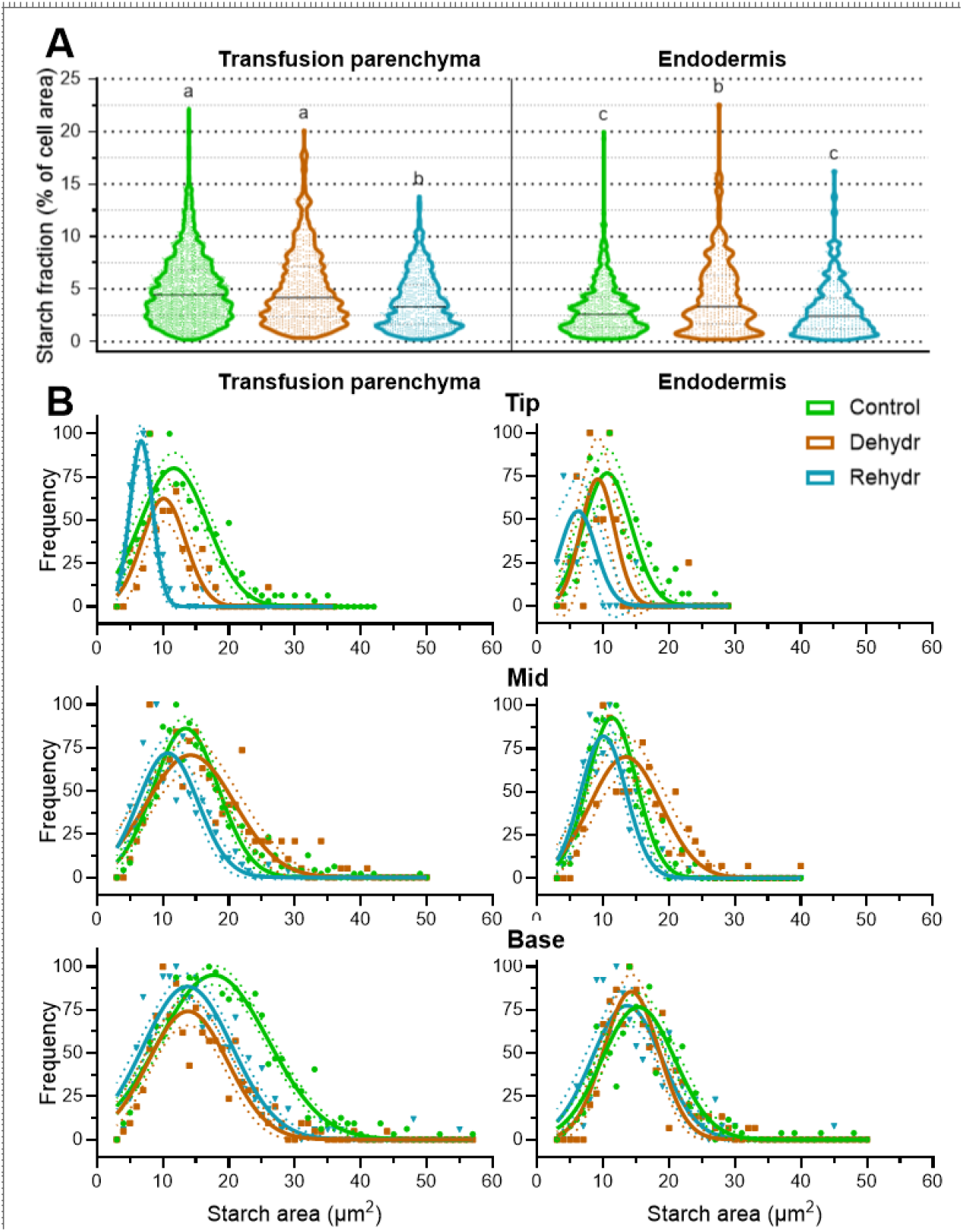
Dehydration-related starch changes in transfusion parenchyma and endodermis. **A:** Starch fraction of tp and en cross-sectional cell areas integrated over tip, mid and base segments of tomographic microscopy series, each segmented at three positions (tp: n>509 per treatment; en: n>278 per treatment). The starch fraction of tp cells decreased and was smallest after rehydration. Due to their decrease in volume (Fig. 5E), the starch fraction of en cells appease to be larger after dehydration but resumed the control value after rehydration. Different letters mark significant differences (*P*<0.0001 according to one-way ANOVA followed by Tukey’s post-hoc test). **B:** Normalized frequency distributions of starch grain sizes at tip, mid and base needle section for tp and en. Curve fittings (lines) assume a Gauss distribution of the grain sizes. Dehydration followed by rehydration reduced the cross-sectional starch area in tp and en cells of the tip and mid-section, as indicated by the frequency of smaller areas. By contrast, smaller grain sizes were in tp of the base section already found after dehydration but there was no further reduction after rehydration. In en cells of the base section, starch grains did not change their size. Confidence intervals at the 95% level are indicated.

Interestingly, 24 h after detachment (before rehydration) we observed positional effects along the needle. In the tip segment, grains were already reduced in size, while in the mid segment they even seemed to be larger than controls. By contrast, in the base segment, starch grains in tp cells reached their final reduction already before rehydration, while those in the en layer showed generally very small changes (Fig. 6B). A stop in phloem export leading to assimilates piling up in the basal parts of the needle - like that observed using esculin (Gao *et al*., 2023) - might be responsible for these positional effects..

## Discussion

Synchrotron imaging using phase-contrast tomography enabled us to visualise and quantify the transfusion tissue deep inside living needles of *P. pinea*. Even the number and size of starch grains could be assessed in tp and en cells; however, phase-contrast typical halo artefacts and a generally smaller size made transitional starch in mesophyll cells inaccessible. Important physiological factors could be studied such as cell size changes along the needle and during a 24-h day cycle, concomitant changes of the cellular starch fraction and adaptations of the volume of the sponge-like tt system.

Importantly, water and nutrients leaving the axial xylem tracheids are bound to the tt system to reach the endodermis, the plasma membrane of which they must pass since Casparian strips block any further apoplasmic transport. Opposing this transport, assimilates entering the en layer from the mesophyll can only move on symplasmically through tp cells that form tortuous symplasmic pathways to Strasburger cells at the phloem flanks (Gao *et al*., 2023; Mai *et al*., 2024). Starch grains evident in cells of the stele (Fig. 1) can only originate from the inwards stream of assimilates through tp cells (Vollenweider *et al*., 2026); photosynthesis in these cells would at the most be marginal, since the en layer excludes any considerable CO_2_ import into the stele. Accordingly, tp cells’ starch metabolism might be more similar to that for storage starch than transitory starch (Lloyd and Kossmann, 2015). However, the observed diurnal fluctuations in the starch fraction contradict a long-time storage function of tp starch.

### Starch content and cell volumes reflect different osmotic conditions along the needle

An unexpected result was the uphill gradient in the cellular starch fraction from tip to base of the long *P. pinea* needles (Fig. 2). The starch fraction is a physiologically relevant indicator being independent of cell size and needle dimensions. The starch gradient was unexpected, since the symmetrical shape of pine needles and the radial organisation of the mesophyll together with the regular spacing of stomata would rather imply an even distribution of photosynthesis, and thus, an even production of assimilates along the needle axis. At the same time, the phloem export from long needles, as those of *P. pinea*, is challenged because of their linear architecture. We have shown before that efficient phloem export from the tip part of needles longer than 60 mm is anatomically realized by grouping sieve elements and allowing phloem loading only from the outermost sieve elements groups, which are in contact with Strasburger cells at the phloem flanks (Fig. 1; (Han *et al*., 2019; Liesche *et al*., 2021; Rademaker *et al*., 2017). This prevents sugar stagnation and accumulation in the tip. According to our present data, starch instead builds up in the base, indicating piling up of photoassimilates in the pre-phloem pathway when meeting the axial phloem stream from the upper needle segments.

In the needle stele, the osmotic conditions are determined by two factors: the cytoplasmic carbohydrate concentrations in tp cells and the water potential in the tt apoplast, i.e., the space contacting each single tp cell and the inner surface of en cells. It is well established that plant cells can osmoregulate under abiotic stress conditions by balancing their mono- and disaccharide levels against starch synthesis or degradation (Dong and Beckles, 2019; MacNeill *et al*., 2017; Thalmann and Santelia, 2017). Any biochemical or molecular characterisation of starch-forming and degrading enzymes is at present not available for conifer tp or en cells (but see (Vollenweider *et al*., 2026).

However, it is safe to postulate that the signalling sugar threhalose-6-phosphate also in conifers is involved in controlling starch fluctuations, as shown in a phylogenetic analysis of the phosphatase family degrading trehalose-6-phosphate (Kerbler *et al*., 2023). Synthesis and degradation of trehalose-6-P lead to formation and degradation of starch, respectively (Annunziata *et al*., 2025; Kolbe *et al*., 2005; Liu *et al*., 2025).

The osmolality of starch molecules is negligible in comparison to that of the glucose molecules or disaccharides like maltose and sucrose they are readily converted to. A smaller starch fraction in tp cells of the tip and mid segments, compared with that of the base, indicates a higher rate of starch mobilisation, i.e. degradation to di- and monosaccharides under the assumption that the concentration of photoassimilate input is constant along the needle. This leads to a higher rate of water endosmosis into tp cells of the upper segments. Thus, the tt apoplast achieves a more negative water potential and will pull up water from the xylem.

The linear architecture of long needles poses the same challenge for xylem unloading as described for phloem loading. The driving force for water into the needle is tension; values might amount to less than –2 MPa in the xylem (Arend *et al*., 2021; Jensen *et al*., 2016). Definite values for pressures differences along needles are unfortunately not available; a paper accounting for the entire pathway from root to needle places the ultimately lowest pressure at the air/liquid interface on the mesophyll cell walls; here approaching down to minus 40 MPa (Lechthaler *et al*., 2020).

How the hydraulic system of xylem tracheids is optimised in the axial direction was discussed in early modelling work on the hydraulics in single vein leaves and pine needles. The authors found that the needle is optimised in its length by decreasing its conductive area from base to tip (Zwieniecki *et al*., 2006). They showed this with experimental data on number (decreasing) and radius (unchanged) of tracheids in *P. sylvestris*, as also confirmed for *P. pinaster* (Bicego *et al*., 2025; Gao *et al*., 2023). To ensure xylem transport to the tip of long needles, osmotically active assimilates in the pre-phloem pathway contribute to the pulling force, mainly provided by transpiration via stomata across the endodermis. This contribution might be small during the day but could well play a major role in maintaining the tension in the apoplasmic tt space extending the axial xylem. Given the same rate of photosynthesis and export, a greater demand for osmotic pressure in the upper segments would then leave less sugar for starch synthesis, as we have observed here.

### At night the driving force for water ascent switches from transpiration to osmotically generated xylem tension as suggested by diurnal starch and volume changes in the needle

The formation of transitory starch in angiosperm leaves starts from zero in the morning, i.e., enzymes involved in the initiation of starch granule formation are essential for transitory starch in mesophyll chloroplasts (Muntaha *et al*., 2022). By contrast, we did not observe starch-free tp or en cells in our study. However, we did observe fluctuations of the starch fraction in tp cells over the day (Fig. 3). Taking changes in grain area as a proxy for starch formation or degradation, we consider the starch, observed in tp cells, to be diurnally manageable. This is supported by the finding that in tip and mid needle the smallest starch grains were found at the last night timepoint, i.e., before onset of photosynthesis, indicating that most of the starch was degraded at this timepoint (Fig. 3).

Our osmoregulation hypothesis was tested by assessing diurnal changes in the cross-sectional cell areas, representing volume changes of the relevant cells and whole tissues. Fluctuations of tp and en cells volumes were moderate, with the significantly smallest volumes in the early afternoon. By contrast, the apoplasmic space of the tt system showed large volume changes over the day, with the smallest volume at the last day timepoint and largest volume at the first night timepoint (Fig. 4A).

The tt system acts as a continuous apoplasmic system since each of its tracheids is connected by conifer-type bordered pits that allow free water exchange. We showed extreme volume changes of nearly 50% in the tt space, enabled by the thin and foldable cell walls between the individual tracheids (see Fig. 1). Volume changes in that scale can only be brought about by osmotic water uptake and release. Water can (i) either come from the axial xylem and leave over the en cell layer, and/or (ii) be exchanged between tt, tp and en.

Ad (i): Water imported in needles through the axial xylem is distributed radially by the tt system to enter the en cells. Aquaporins allowing entry into and exit from en cells have been documented in pine needles as well as in their tp cells (Laur and Hacke, 2014). How these are actively regulated to control the water relations across the endodermis is unclear and not even solved in case of the well-studied roots (Couvreur *et al*., 2018; Lechthaler *et al*., 2020). Exit from the endodermis to the mesophyll will strongly depend on the transpiration rate and, thus, on stomatal opening. In a related study, we measured for *P. pinea* a strongly reduced water evaporation rate in needles kept in the absence of light (Gao *et al*., 2023). Accordingly, the pulling force of stomatal evaporation on the outside of the en cell layer will be much less at night than during the day.

Ad (ii): Changes in tissue proportions give an indication of the direction of water exchange (Fig. 4B), or rather give us a snapshot of what had happened at the indicated timepoints.: water went from tp to tt from last night to afternoon timepoint, from tt to both tp and en at the last day timepoint, and from en to tt at the first night timepoint. This timepoint is characterised by the largest swelling of the tissue inside the endodermis in all three needle segments (see Supplementary Fig. S4). Since evaporation is smallest at night, the swelling of the vascular tissue is compatible with the hypothesis that water is osmotically pulled up in the needle. To pull it up to the upper segments, tp cells in these needle segments need to invest more in starch degradation than in mid and base, and, therefore have a smaller excess of carbohydrates available to build transitional starch (Fig. 3).

The question remains, how starch metabolism and turgor changes are coupled. The turgor of living tp cells is depending on cell wall properties (tensile strength of cell walls) and the osmotic balance between tp and tt. Sugars arriving in the pre-phloem pathway will lead to turgor increase but also to a (delayed?) switch of starch production. A role in these dependencies might be mechanosensitive plasma membrane receptors that respond to turgor changes. Relevant mechanosensitive calcium channels, involved in the regulation of cell turgor, have been characterised in Arabidopsis and are present in all land plants, also gymnosperms (Ali *et al*., 2023; Nishii *et al*., 2021; Yoshimura *et al*., 2021).

### Osmotically-generated tension in the xylem contributes to cavitation repair

To test feasibility of the above hypothesis we challenged xylem transport by studying rehydration of detached needles. We expected that detachment and risk of embolies in axial tracheids would render water transport into needles impossible. Detaching the needle means that phloem transport cannot continue, as shown with a phloem tracer that only moved from tip to base of needles when still attached to the tree (Gao *et al*., 2023). Thus, our experiments are not entirely comparable to situations in nature where needles resume from drought-stress and dehydration as a natural event of their climatic challenges (Bowling *et al*., 2018; Maruta *et al*., 2020). Comparing angiosperm and gymnosperm leaves, the specific architecture of the latter with the extended transfusion tissue and Casparian strips might explain the broad climatic adaptations of gymnosperms, considering the outside-xylem vulnerability of angiosperm leaves (Scoffoni *et al*., 2017). Simple detachment as used here is a much more moderate impact on xylem transport in needles than experimental vacuum, drying or freezing treatments (Bouche *et al*., 2016; Cochard *et al*., 2004; Feng *et al*., 2021; Roden *et al*., 2009).

Dehydration led to moderate volume reductions of tp and en cells but to a dramatic shrivelling up of the tt system, including the entire stele. To our surprise, these needles resumed completely, when their base was placed in water, regarding both size of the living cell types tp and en and proportion between those and the apoplasmic tt system (Fig. 5). The rehydration process was accompanied by degradation of the transitional starch in tp cells, while en cells recovered the starch fraction of the controls. Decrease of starch grain size and starch fraction in tp cells suggests that these cells still are viable some 30 h after detachment as indicated by the observed reduction of the grain size in needles after dehydration-rehydration. The pre-phloem transport of assimilates between en and axial phloem demands a standing assimilate gradient between the production and export sites. This gradient is certainly disturbed in detached needles, where phloem export is arrested. However, *P. pinaster* needles still attached on a tree branch shrunk similarly and resumed thereafter – as shown also with synchrotron tomography by (Bouche *et al*., 2016). Their experiments desiccated the needles using very negative water potential values in quite short time, exposing them to more stressful conditions than our experimental setup and showing – in contrast to our results – that many axial tracheids were filled with air. Under these conditions, the entire needle was drastically deformed. Our dehydration experiments equally led to a deformation of the needle and a moderate reduction of the stelar cross-sectional area (Supplementary Figure S4). As mechanism for resumption of xylem transport, our results suggest that stored starch grains in tp provide a buffer to maintain an assimilate gradient through periods of drought and provide the osmotic potential to resume water ascent in needles.

In Figure 7 we try to summarise the diurnal and dehydration-related volume and starch changes. According to our hypothesis, the plastids in tp cells are filled with starch during the day by using photoassimilates arriving from the endodermis after start of photosynthesis. Starch is then degraded during the night to provide the osmotically generated pulling force for water import into the needle. This force is so strong that it leads to a swelling of the entire stele (Supplementary Fig. S4). Only little water will move on to the mesophyll from the outside of the endodermis, since needles with closed stomata have a very high humidity in the intercellular space of the mesophyll. Moreover, the nighttime stomatal conductance in Pinaceae is very small according to (Yu *et al*., 2019). Our data show that swelling declined already at the last night timepoint, when the starch was used up (Fig. 3; Supplementary Figure S4). Contrasting swelling of the entire stele, dehydration leads to dramatic shrinking of the tt space, and moderate volume loss only of tp and en. Starch is used for the formation of mono- and disaccharides, which maintain water intake into these cells (Fig. 7).

**Figure 7:**
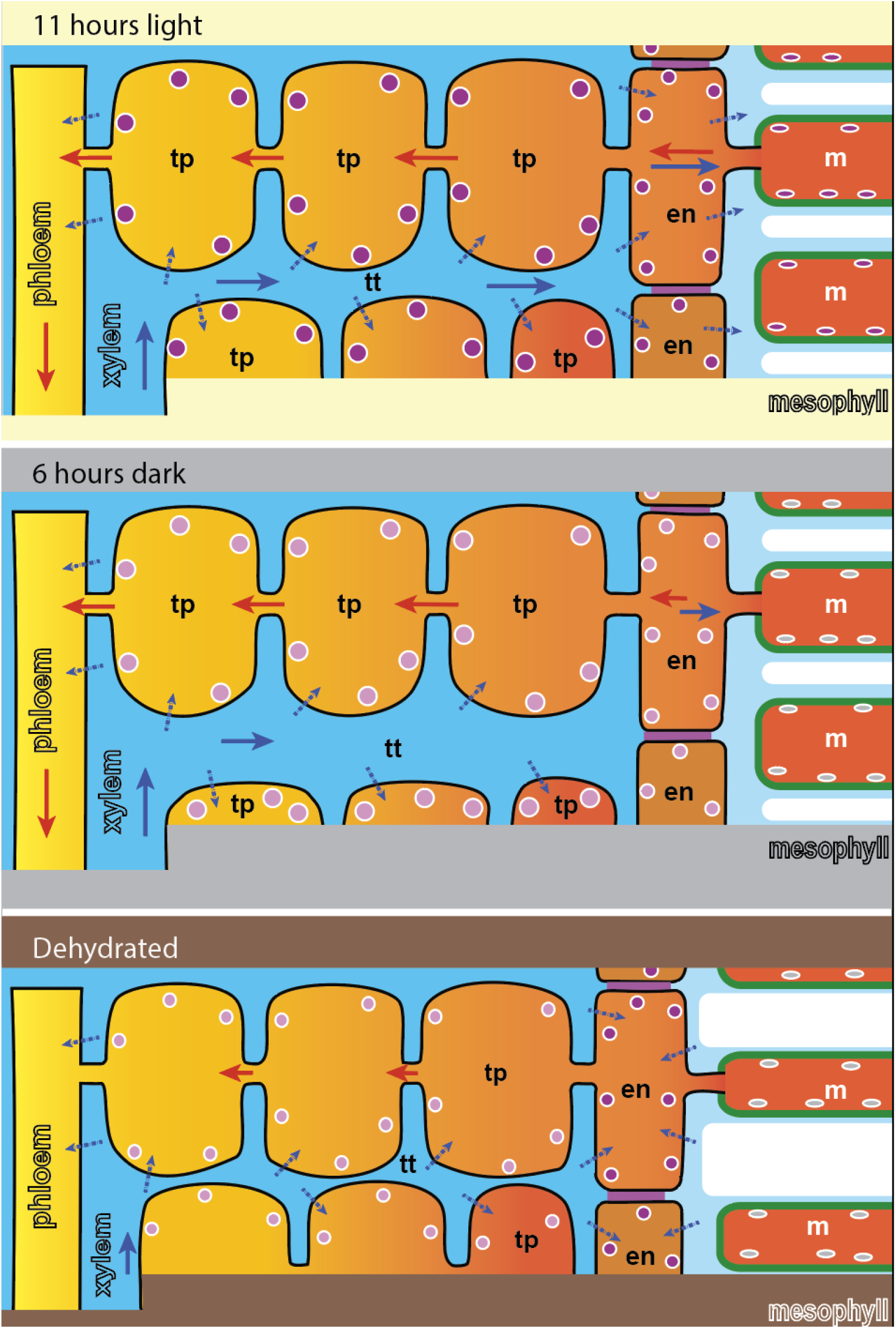
Model of the two most contrasting settings in the diurnal cycle, and of dehydration: Eleven hours light: Excess photoassimilates are stored in starch (violet organelles) of mesophyll (m), endodermis (en) and transfusion parenchyma cells (tp). Blue stippled arrows show water endosmosis via aquaporins into tp and en, driven by a high solute potential in the pre-phloem pathway. Water passes the endodermis layer towards the mesophyll, driven by transpiration. This diurnal stage leads to the smallest volume of the apoplasmic tt space. Six hours without light and photosynthesis: Most of the starch is mobilised (light-violet organelles) and mono- and disaccharides in tp drive endosmosis into tp, pulling up water in the apoplasmic space, thus maintaining ascent of water in the xylem. Since stomata are closed and transpiration negligible, water is not passing the endodermis towards the mesophyll, and tt and tp together achieve their largest volume. Red arrows: assimilate transport, blue arrows water and nutrient transport. Dehydration: Starch content in tp is (light violet organelles) is further reduced. The entire stele is shrunken due to water loss, most of which from the tt space. Assimilate content in tp and en still are turgescent due to their osmotic potential.

Rehydration is possible by resumption of xylem transport, driven by osmotic water intake into tp cells through mobilisation of even more starch in tp cells, resulting in the smallest starch fractions and grain areas (Fig. 6).

## Conclusion

The three main observations in our study – the longitudinal starch gradient, the diurnal fluctuation of starch content and cell volumes, and the resumption of cell and apoplast volumes accompanied with the degradation of starch volumes in rehydrated needles – are strong indications for an active role of tp cells in the recovery of xylem transport and, thus, also for an important role in cavitation repair. Open questions are, how starch formation and degradation are regulated in tp cells. The important setpoints are a threshold value for the lower assimilate concentration, where starch has to be degraded to yield more osmolytes, and the upper assimilate concentration, where an excess of assimilates is removed from the cytoplasm by starch synthesis in tp plastids. These setpoints have at the same time a strong influence on maintaining the assimilate gradient in the pre-phloem pathway from endodermis to the Strasburger cells at the flank of the axial phloem (Mai *et al*., 2024; Vollenweider *et al*., 2026). By sensing small turgor differentials, an individual tp cell might be able to moderate (steepen or flatten) this gradient depending on the number of cells bridging the gap between endodermis and phloem flank.

## Supplementary data

The following supplementary data are available at JXB online:

Fig. S1. X-ray micrographs of vascular tissues in Pinus pinea needles detached immediately before imaging

Fig. S2. Diurnal changes in starch fraction and grains cross-sectional areas at the needle positions, tip, mid and base.

Fig. S3. Diurnal tissue area changes in transfusion tissue and endodermis.

Fig. S4. Changes of the summed area of transfusion parenchyma, transfusion tracheid system and endodermis layer

Fig. S5. Dehydration-related changes in tip, mid and base of P. pinea needles.

Video S1. Animation following all cross-sections through a tomographic microscopy stack of a *P. pinea* needle comprising 1000 transvers slices of 0.325 µm thickness.

## Acknowledgments

We thank the Center for Advanced Bioimaging, University of Copenhagen, for providing its electron and fluorescence microscopes.

## Author contributions

AS and B designed and supervised this study; MS developed and supervised the synchrotron settings; CG, PB and AS prepared the plant material and controlled the PSI’s Tomcat beamline during imaging; CG and AS fixed plant material for electron microscopy; PB and ETH segmented the synchrotron tomographic data; AS drafted the manuscript, analysed the quantitative data and prepared the Figures and videos; all authors contributed to discussion and interpretation of the data.

## Data availability

The data supporting the findings of this study are available from the corresponding author, Alexander Schulz, upon request.

