## Supplementary data for "Stelar starch management tailors diurnal and rehydration-related water flows in *Pinus pinea* needles"

*Contributed equally to this work

### Supplementary data

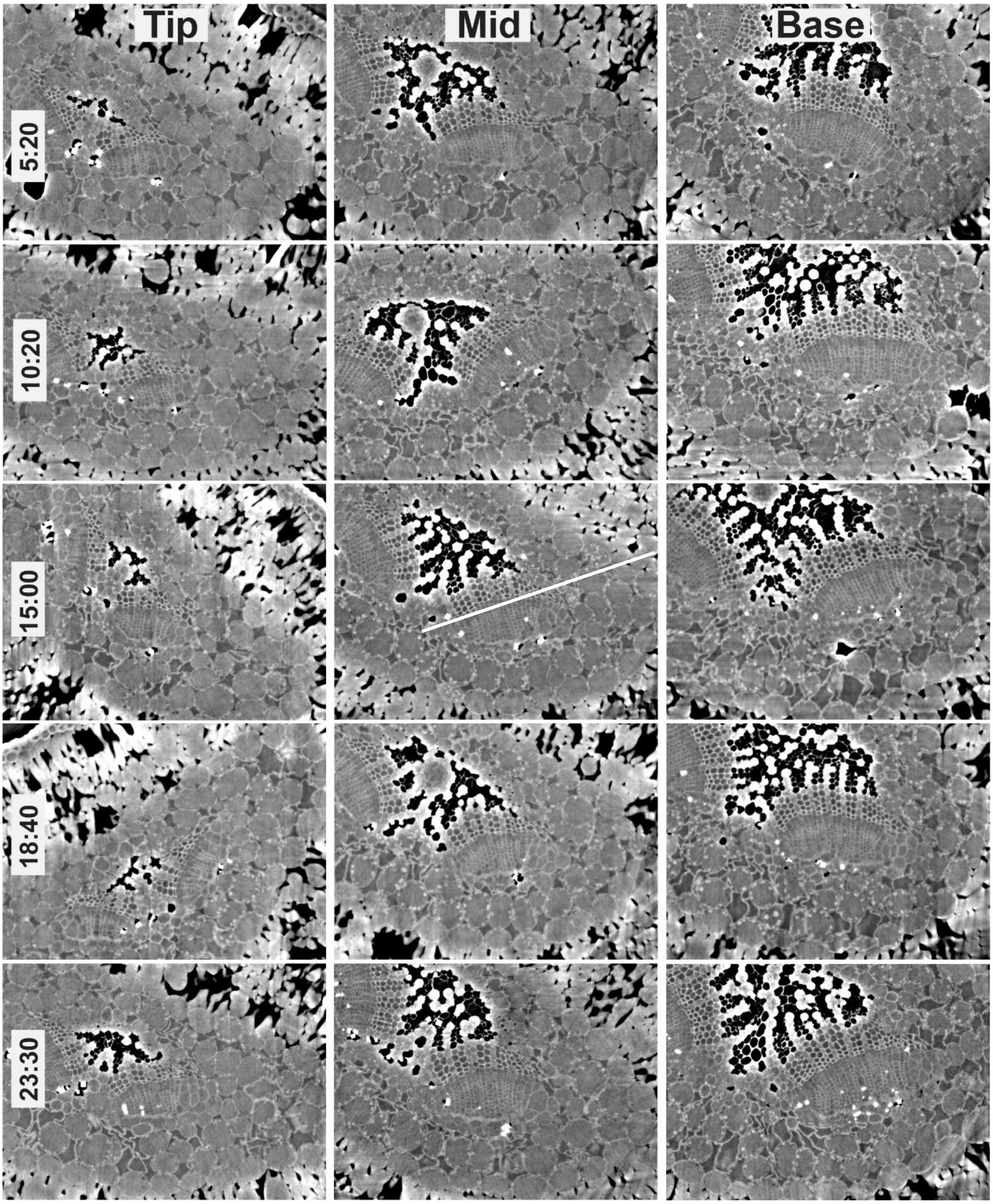


**Supplementary Figure S1: X-ray micrographs of vascular tissues in *Pinus pinea* needles detached immediately before imaging. A:** Representative cross section through a needle at three positions of the needle, the tip, mid and base segment, reconstructed from µX-ray CT series. The transfusion tissue is placed between endodermis and the axial vascular tissue (see Fig. 1A. in the main paper). The line in the 15:00 mid panel shows the location of the longitudinal slice in Fig. 1B. **B:** Tangential view of the mid segment at 15:00 from Supplementary Fig. S1A showing the phloem and the neighbouring transfusion tissue with tp cells containing starch grains (white dots) and the apoplasmic tt space (dark). The transfusion tissue is bordered by the endodermis layer on the right side. Mesophyll cells with intercellular spaces are on the right picture border

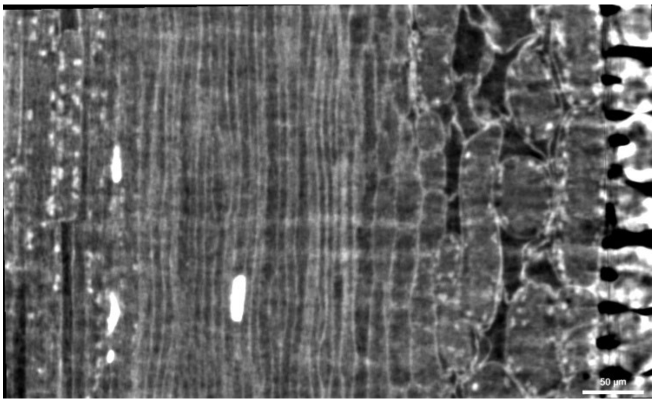

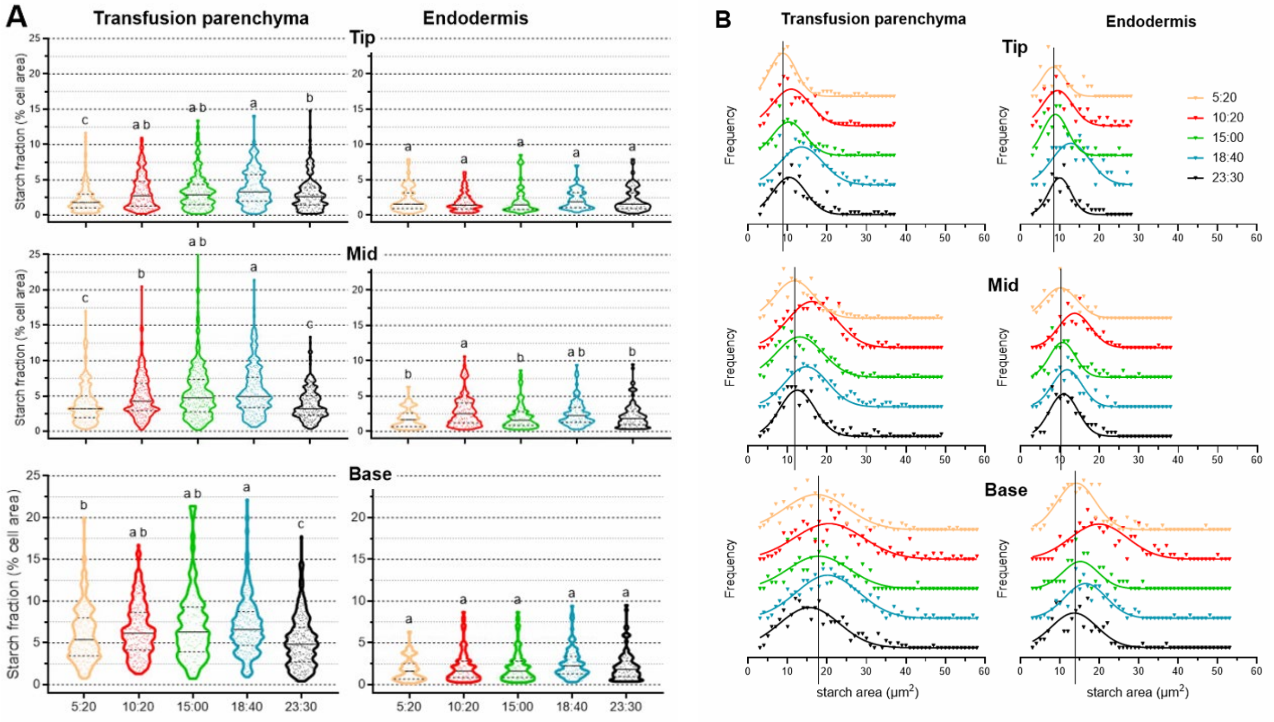


**Supplementary Figure S2: Diurnal changes in starch fraction and grains cross-sectional areas at the needle positions, tip, mid and base. A: S**tarch fraction of tp cross-sectional cell areas in tip, mid and base segments of µX-ray CT series. Different letters mark significant differences (*P*<0.005 according to one-way ANOVA followed by Tukey's post-hoc test).

**B: Normalized frequency distributions of starch grain sizes at tip, mid and base needle section for tp and en.** Curve fittings (lines) assume a Gauss distribution of the grain sizes. The vertical lines indicate the mean values at the last night time point at 5:40. They are smallest in the tip and mid segments of both tissues, but not in the base segment of tp and en cells that have smallest means at midnight. For statistic tables see supplementary data.

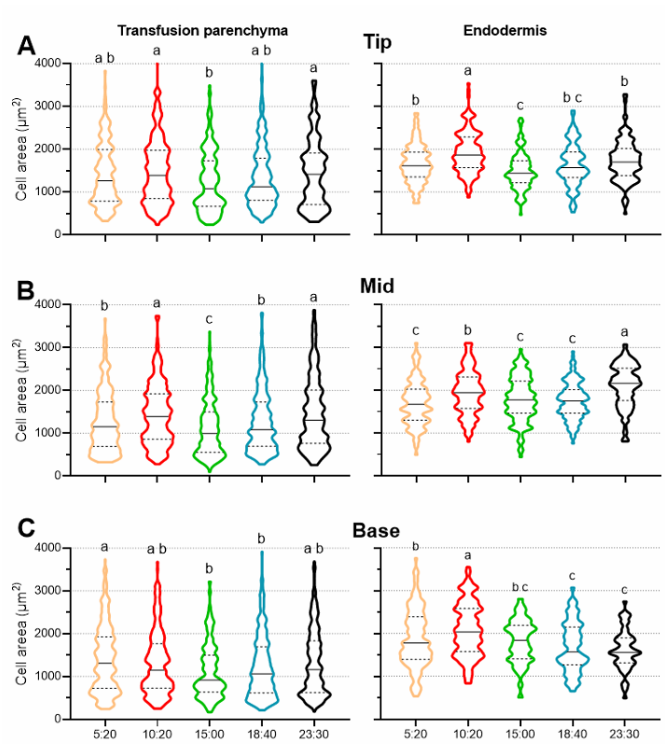


**Supplementary Figure S3: Diurnal tissue area changes in transfusion tissue and endodermis. A, B:** The cross-sectional cell sizes of tp and en in tip and mid segment were smallest in the afternoon and increased up to midnight (tp n mean >236; P<0.002); en mean=152;P<0001). (tp n average = 323>; P<0.0001); en n average=159; P<0001). **C:** The cross-sectional cell sizes of tp in the base segment decreased from morning to afternoon and increased again up to midnight (n >125; P<0.006); Cross-sectional area of en cells was highest in the morning and decreased thereafter (n=120;P<0001).

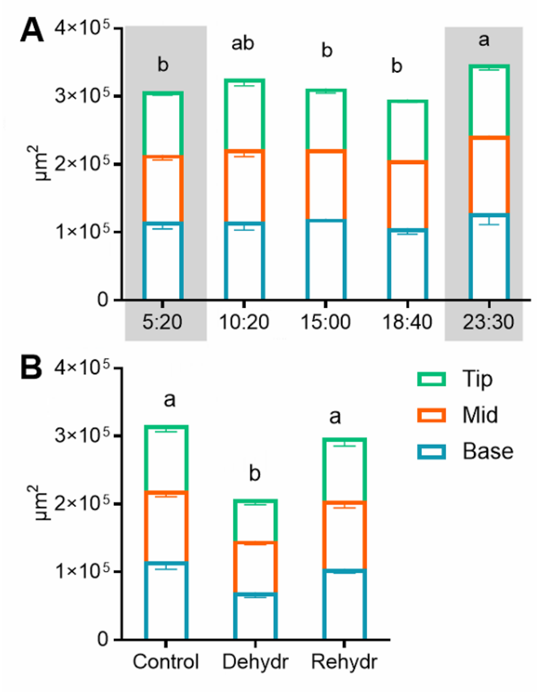


**Supplementary Figure S4: Changes of the summed area of transfusion parenchyma, transfusion tracheid system and endodermis layer. A, Diurnal changes:** At all time points, the tip area was smallest, and the base largest. The summed area did not change significantly between last night and last day timepoint. However, all needle postions were largest at midnight. **B, Dehydration-related changes**: The summed area was strongly reduced under dehydration and resumed fully after rehydration, except for the base segment. The area difference between tip, mid and base segments was equilibrated under dehydration. Letters assign significant differences according to one-way ANOVA followed by Tukey's post-hoc test (P<0.0001)

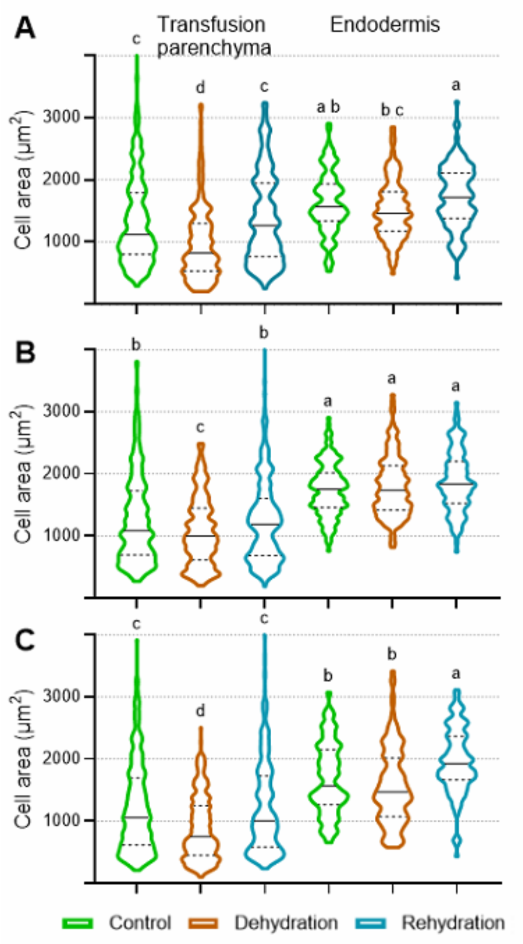


**Supplementary Figure S5: Dehydration-related changes in tip, mid and base of P. pinea needles.** The cross-sectional cell area of transfusion parenchyma (tp) cells drops due to dehydration and resumes its control values after rehydration. By contrast, the cell area of endodermis (en) cells does not change under dehydration in all needle segments but increases in tip and mid segment after rehydration. Different letter assign significant differences according to one-way ANOVA followed by Tukey's post-hoc test (*P<0.0001*; tip: n=> 188 for tp, >115 for en; mid: n=> 286 for tp, >138 for en; base: n=> 331 for tp, >149 for en).
